# Exposure to non-nestmate odours changes the odorant receptor profile in *Acromyrmex echinatior* ants

**DOI:** 10.1101/2025.08.17.668880

**Authors:** Mélanie Bey, Naomi Alex, Lisa Maczkowicz, Volker Nehring

## Abstract

Insects depend on a broad olfactory perception ability that involves many sensory receptors. Social insects, in particular, use olfactory cues to maintain colony cohesion (Lenoir et al., 1999; Leonhardt et al., 2016). They recognize nestmates through colony-specific olfactory labels that individuals store as neural templates in their memory (Sherman et al., 1997). Learning continuously optimises the nestmate recognition template to keep up with the constant changes in colony labels (d’Ettorre and Lenoir, 2010; Errard and Hefetz, 1997; Wilgenburg et al., 2012). The template is often considered to be located in higher brain centres and potentially a product of learning (Bey et al., 2025; Brandstaetter et al., 2011; Esponda and Gordon, 2015; van Zweden and d’Ettorre, 2010). However, some authors suggest it might be in the neural periphery, i.e. the antennae or antennal lobes, formed by habituation or receptor adaptation (Guerrieri et al., 2009; Ozaki and Hefetz, 2014; Stroeymeyt et al., 2010). Here we investigate a potential mechanism for the construction and functioning of the colony template in the peripheral nervous system: the antenna’s odorant receptor (OR) profile and its dynamics. We exposed *Acromyrmex echinatior* leaf-cutting ants to non-nestmate odours and analysed the consequences on their behaviour and antennal gene expression. Consistent with other studies, prolonged exposure to non-nestmate odours reduced worker aggression towards the non-nestmate label, indicating habituation to the non-nestmate odour (Carlin and Hölldobler, 1983; Errard and Hefetz, 1997; Stroeymeyt et al., 2010). Exposure also altered the expression of the OR genes. Notably, the OR profiles were colony-specific, mirroring the colony-specific recognition labels. When we exposed two different colonies to the same non-nestmate odour, the colony-specificity of the odorant receptor gene expression vanished. This indicates that the olfactory machinery used to perceive nestmate recognition cues is flexible and adapts to the current nest-specific olfactory environment. The OR profiles could either become more sensitive to the nestmate recognition cues by increasing the number or ORs for the nest-specific substances, or less sensitive by decreasing the expression of these ORs. The latter would in turn increase the sensitivity to non-nestmate cues. This would be a mechanism explaining habituation to nestmate recognition cues that has been described across the social insect literature.

## Introduction

Every species perceives the world in its own way. Evolution created notable examples of specialised perception such as the trichromatic vision in primates (Tan and Wen-Hsiung, 1999), the remarkable somatosensory organs of the star-nosed mole (Marasco and Catania, 2007), and the fascinating olfaction abilities of the insects (Wyatt, 2014). Insects require perception of a large repertoire of pheromones and other odorants in order to perform species-and mate-recognition, or simply to forage successfully. Insects perceive reality beyond the capacity of the human mind to comprehend (Benton, 2022) and employ a multitude of sensory receptors, including odorant receptors (ORs). Social insects rely on olfaction to distinguish between nestmates and non-nestmates, a crucial capacity that allows division of labour and provides protection against intruders and parasites (Leonhardt et al., 2016; Sturgis and Gordon, 2012; van Zweden and d’Ettorre, 2010).

Nestmate recognition is based on the identification of colony-specific odours (“labels”) that insects bear on their cuticles. The labels are composed of a cuticular hydrocarbon mixture, which workers compare to a neural template of their colony’s olfactory identity (Sherman et al., 1997). The template can change over time when the colony label changes (Errard and Hefetz, 1997; Le Moli and Mori, 1990; Lenoir et al., 2001; Leonhardt et al., 2007). Numerous studies investigated the behavioural aspects of nestmate template dynamics and showed that social insects exposed to non-nestmate odour integrate the non-nestmate labels into their templates, so that they then treat the former strangers as if they were nestmates (Carlin and Hölldobler, 1983; Dahbi et al., 1996; Foubert and Nowbahari, 2008; Guerrieri et al., 2009; Neupert et al., 2018; Stroeymeyt et al., 2010). While the behavioural effects are well described, the underlying physiological mechanisms remain elusive. It is unclear whether the template construction and its flexibility is based on sensory adaptation, habituation, or another form of learning. We know that social interactions with nestmates such as trophallaxis or allogrooming (Dahbi et al., 1999), as well as some forms of associative learning (Bey et al., 2025; Bos and d’Ettorre, 2012), are required for template construction and modification. The nature of the template and its location in the nervous system is subject to ongoing debate, giving rise to two hypotheses: memory formation in the central nervous system (Brandstaetter et al., 2011; van Zweden and d’Ettorre, 2010) versus peripheral adjustments in recognition (Guerrieri et al., 2009; Nehring et al., 2016; Ozaki and Hefetz, 2014; Stroeymeyt et al., 2010).

The memory formation hypothesis is based on associative learning, the association of colony odours with experiences individuals make with members of these colonies (e.g. attacks, grooming). This type of learning easily explains the variation in recognition templates and recognition skills observed across workers (Bey et al., 2025; Esponda and Gordon, 2015; Larsen et al., 2016, 2014; Norman et al., 2014).

The theory behind the peripheral recognition mechanism suggests that individuals cease to react to the nestmate labels that they are constantly exposed to (d’Ettorre and Lenoir, 2010; Ozaki and Hefetz, 2014; Reeve, 1989; Wilgenburg et al., 2012). They will only react to novel stimuli, which would include non-nestmate labels. There are two mechanisms potentially involved in this process: habituation and sensory adaptation. Habituation is defined as a decrease in the behavioural response to a constant stimulus (Rankin et al., 2009) and is a simple form of non-associative learning (Kandel, 2004). Over time, social insects might gradually ignore their own colony odour and become more sensitive to other odours, e.g. those of non-nestmates. A series of experiments showed that the process of habituation appears to necessitate neurophysiological changes such as synapse plasticity, leading neurons to stop transmitting the information e.g. in the antennal lobe (Guerrieri et al., 2009; Langen et al., 2000; Stroeymeyt et al., 2010). Sensory adaptation, in contrast, is a modulation of the olfactory receptors (ORs) at the dendritic membrane of the olfactory sensory neurons in response to a prolonged exposure to a new olfactory environment (Todd and Baker, 1999; Wark et al., 2007). ORs that are overstimulated by specific odorants undergo a process of desensitization, resulting in the loss of perception of the odorants (d’Ettorre and Lenoir, 2010; Wilgenburg et al., 2012). Conceptually, both habituation and sensory adaptation blind out stimuli that are invariably present, and allow individuals to continuously be able to perceive relevant stimuli (those that still vary) from their environment and to respond appropriately.

In mammals, a molecular mechanism for olfactory optimisation at the OR level has been observed. When exposed to odours, mice modulate their OR gene expression (Ibarra-Soria et al., 2017; Von Der Weid et al., 2015). Similar alterations in mRNA levels were reported in *Drosophila melanogaster* (Koerte et al., 2018; Von Der Weid et al., 2015). In theoretical models it was shown that a modulation of OR gene expression and an assumed change in the relative proportions of different ORs in the olfactory periphery (“OR profiles”) can optimise the perception of important cues. How strong OR expression adjustments need to be depends on receptor tuning: The OR profiles should react more strongly to environmental change when receptor tuning is narrow, and less strongly when there is a large total quantity of neurons (Teşileanu et al., 2019). Insect odorant receptors are G-protein-coupled receptors composed of 3 co-receptor units (ORco) and a single ORx protein that is odorant-specific (Suh et al., 2014; Zhao et al., 2024). This complex acts as a metabotropic ionotropic receptor (Gomez-Diaz et al., 2018). In ants, the ORx subunits are thought to be narrowly tuned to single or few odorants (Pask et al., 2017).

For the specific case of ants, previous studies suggested that the particular ORs that an individual expresses are optimised for the tasks that individual performs. Foragers seem to express a specific trail pheromone receptor (Koch et al., 2013) and their overall OR gene expression profile also differs from that of nurse ants (Caminer et al., 2023; Koch et al., 2013).

Since OR profiles of individuals vary and OR gene expression is known to respond to changes in the olfactory environment, we speculated that dynamic regulation of OR profiles could also be a mechanism for the habituation-type learning of nestmate recognition odours in the olfactory periphery. If individuals produced fewer ORs binding those substances that are typical to the nestmate odour, e.g. by downregulation of OR gene expression, the sensitivity to nestmate odour would decline as it has been suggested to happen in some circumstances (Guerrieri et al., 2009; Ozaki and Hefetz, 2014; Stroeymeyt et al., 2010). In turn, more neurons could be equipped with ORs that are tuned to other substances, which would increase the sensitivity to intruders bearing novel odours.

We show, in line with studies on other species (Carlin and Hölldobler, 1983; Errard and Hefetz, 1997; Guerrieri et al., 2009), that prolonged exposure to non-nestmate odour reduces the aggression of *Acromyrmex echinatior* workers towards the non-nestmate label. We also found that OR gene expression is colony-specific and that it reacts dynamically when individuals are exposed to nestmate recognition labels. The odour exposure affected the OR gene expression profiles in a predictable way, and also affected the expression of other genes that might be involved in regulating OR profiles in the antenna. Interestingly, when we exposed different colonies to the same non-nestmate cuticular hydrocarbon extract, the initial separation between colonies in the OR gene expression profiles disappeared. This indicates that, as their olfactory environments became similar, their OR profiles also became similar, which could indicate that the nestmate recognition template is in part encoded in the odorant receptor profiles.

## Results

Ants are known to incorporate novel odours into their nestmate recognition templates and to habituate to non-nestmate odours during constant exposure. We explored this effect in *Acromyrmex echinatior* ants by exposing workers to non-nestmate cuticular hydrocarbon (CHC) extracts in experimental subcolonies for three days (Fig. 1A). We set up three odour exposure treatments: the control, in which the ants were only exposed to the solvent (pentane); the conspecific exposure, where the ants were exposed to CHC extract of non-nestmates of the same species; and the allospecific exposure, where ants were exposed to the CHCs of another leaf-cutting ant species, *Acromyrmex octospinosus*. We then measured the duration of mandible opening exhibited by ants towards the CHC extract they had been exposed to, and compared it to the control ants’ responses towards the same CHC extract (Fig. 1). When focal ants had previously experienced the allospecific non-nestmate CHC extract, they significantly reduced their aggression towards that odour compared to the control group (Fig. 1C, n = 56, F_1,51_ = 11.93, p < 0.01; Tab. S2). While there was a similar pattern in the conspecific treatment (Fig. 1B), the results were not as conclusive (n = 176, F_1,168_ = 0.15, p = 0.70) because the effect depended on the colony origin of the focal ants (treatment x colony interaction p < 0.05, Tab. S1). Combined with results reported from other studies – the effect of habituation seems to depend on the colony combination (Carlin and Hölldobler, 1983; Couvillon et al., 2007; Guerrieri et al., 2009; Stroeymeyt et al., 2010) this evidence suggests that the nestmate recognition template changes towards the non-nestmate odour after exposure to this odour.

**Figure 1.**
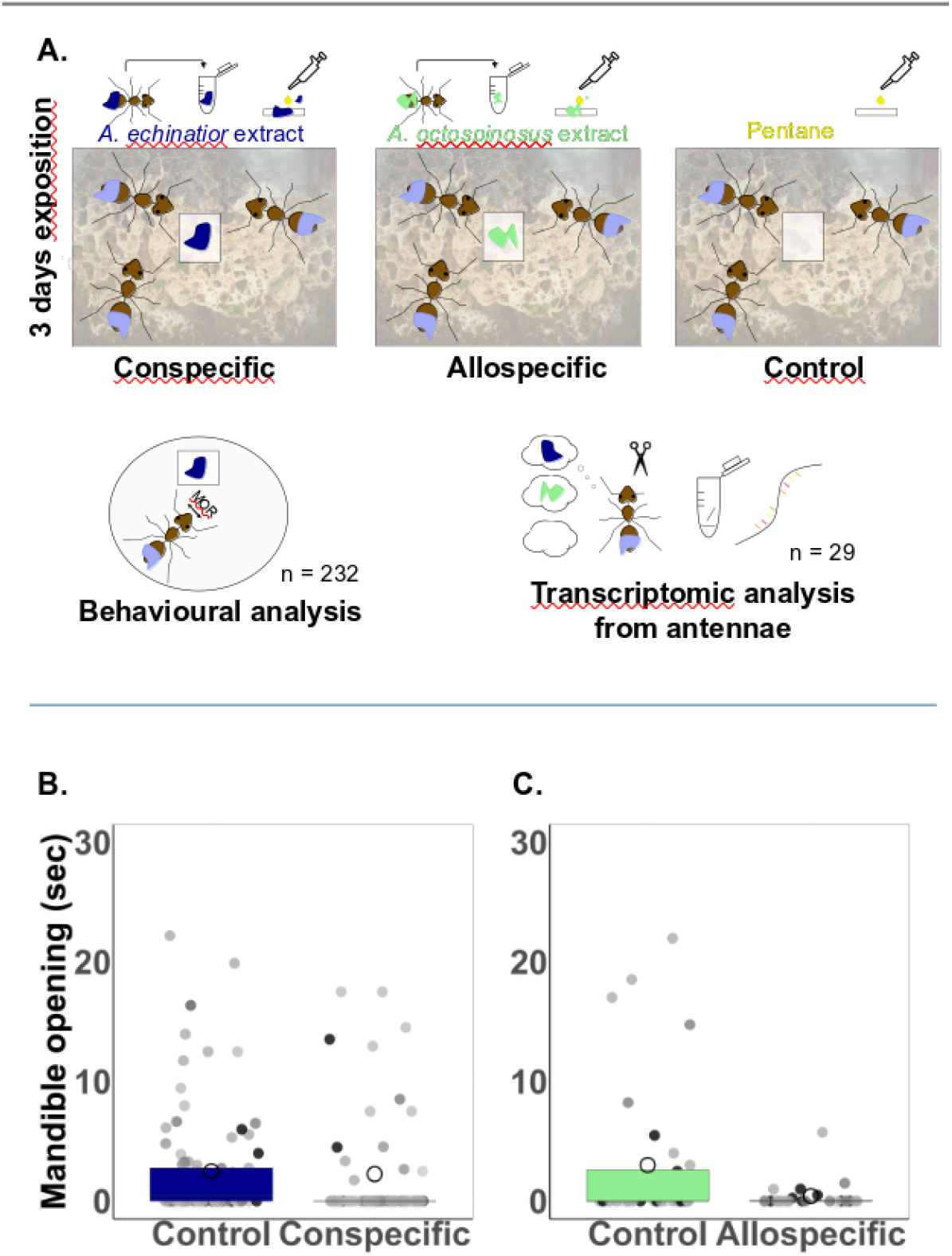
Experimental design and behavioural habituation. A) We established 3 treatments of odour exposure by setting up 42 subcolonies of 8 to 10 ants from 7 *Acromyrmex echinatior* original colonies and continuously exposed them to either non-nestmate CHC extract from A*cromyrmex echinatior* (conspecific), *Acromyrmex octospinosus* (allospecific), or to solvent (control). On the 4th day, we measured the aggression of the ants (mandible opening duration) and extracted RNA from the antennae for transcriptome analysis. The ants that had been previously exposed to the non-nestmate CHC extract generally reduced their aggressive response towards the same CHC extract compared to the control ants, with a more consistent effect for the allospecific exposure (B: conspecific, n = 176, p = 0.70, Tab. S1; C: allospecific, n = 56, p < 0.01, Tab. S2). For the behavioural experiment, twelve different combinations of 7 focal colonies and 8 non-nestmate colonies were used. Each dot is the measurement of an individual ants’ aggression score. The different shades correspond to the colony origin of the focal ants. The large circles indicate arithmetic means and the box plots medians and interquartile ranges.

Some models predict that the nestmate recognition template may already be encoded at the peripheral level in the antenna (Ozaki and Hefetz, 2014). Because it has been suggested that dynamic OR gene expression might optimise the perception of important cues in vertebrates (Ibarra-Soria et al., 2017; Teşileanu et al., 2019), we suspected that the production and hence the overall profile of odorant receptors itself might play a role in template formation in ants as well. We predicted that the profile may change when the recognition template changes, contributing to the habituation effects observed in nestmate recognition behaviour. To test this hypothesis, we analysed the antennal gene expression of ants from two colonies that we exposed to the same non-nestmate CHC extracts (con- and allospecific) or to solvent control, as in the behavioural experiment.

The expression pattern of OR genes in the antennae was colony-specific in the control treatment (Fig. 2A, n = 10, Manova: Pillai’s Trace = 0.95, F_2,6_ = 66.2, p < 0.001). This is expected when OR repertoires are be optimised for the perception of important substances, and hence for nestmate recognition. When we exposed both colonies to the same non-nestmate colony CHC extract, the separation between colonies in the OR gene expression disappeared (conspecific: Fig. 2B, n = 10, Pillai’s Trace = 0.40, F_2,7_ = 2.3, p = 0.17; allospecific: Fig. 2C, n = 9, Pillai’s Trace = 0.10, F_2,6_ = 1.18, p = 0.78). This was confirmed in a Dseq2 analysis where 52 out of 435 OR genes were differentially expressed (p_adj_ < 0.05) between control ants of the two colonies (Tab. S5), but this number was significantly reduced to 6 and 7 OR genes after the exposure to con- and allospecific non-nestmate CHC extract, respectively (Tab. S5; Pearson’s χ^2^-tests on differentially vs. non-differentially expressed genes between the three treatments, χ^2^ = 67.06, df = 2, p < 0.001). Such a pattern indicates that the changes in OR gene expression are specific to the odorants the ants were exposed to. When ants habituate to the CHC extracts of another colony, which effectively changes the nestmate recognition template, changing the OR gene expression might optimise the sensitivity of the antennae to improve perception of the most relevant cues and accelerate the discrimination of nestmates from non-nestmates. Because the expression of OR genes was colony-specific, we analysed separately for both experimental colonies which ORs were affected by which odour treatment as compared to the control. The exposure to conspecific non-nestmate odour affected the expression of 20 OR genes in colony Ae32, and allospecific odour that of 3 genes (p_adj_ < 0.05, Fig. 2D, Tab. S6). In colony Ae21, no OR genes were significantly affected after correcting for multiple testing. However, the overall profile of the OR gene expression always changed as can bee seen from the frayed shape of the gene expression histograms in Fig. 2D, so that OR profile optimisation might not be a matter of strong changes in single ORs but small changes in many.

**Figure 2.**
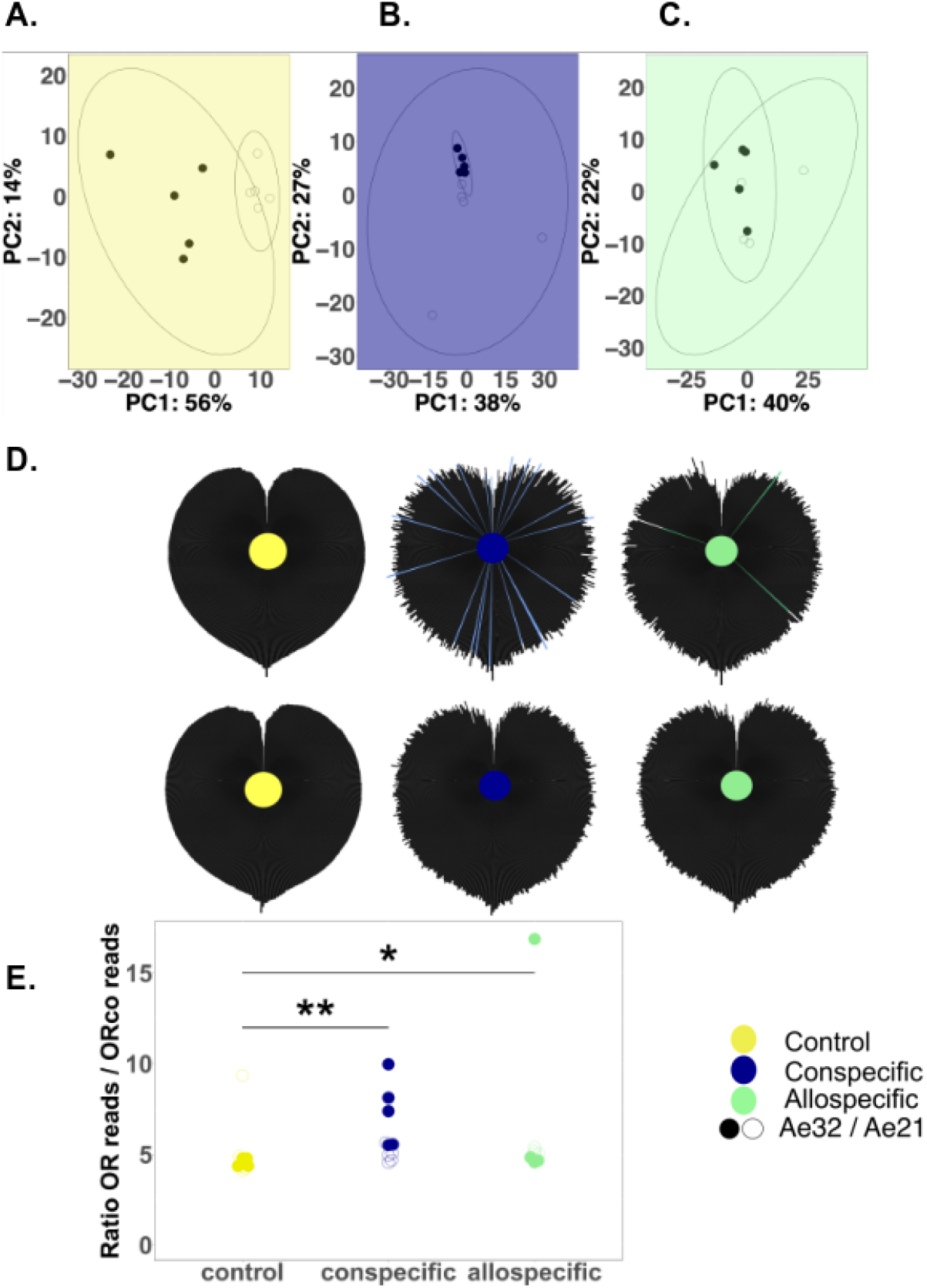
Colony identity and odour exposure affect antennal gene expression. The expression of OR genes expression differed between the two colonies in the control group (A; n = 10, MANOVA p < 0.001). This difference disappeared once both colonies were exposed to the same non-nestmate CHC extract, either from a conspecific colony (B; n = 10, p = 0.17) or an allospecific colony (C; n = 9, p = 0.78). Each dot represents an individual ants’ OR gene expression. Filled and empty dots indicate the two different colony origins of the focal ants. Exposing ants to non-nestmate odours changed their OR gene expression profiles (D). Exposure to CHC extracts from conspecific (blue) or allospecific (green) non-nestmates modulated the OR profiles in the focal ants’ antennae. Each of the 434 bars displayed around the colour dots indicates the average relative expression level of one OR gene. In all three subfigures, bars are displayed in the same order, which is sorted according to their expression levels in the control (left/yellow). Genes with the lowest relative expression are on the top, and those with the highest expression on the bottom of the leaf shape. Bars for OR genes that were differentially expressed compared to the control (Dseq2 p_adj_ < 0.05) are highlighted in colour. The smooth shape of the control’s OR profile becomes frayed in the treated groups, indicating that the overall profile changed. Exposing ants to non-nestmate odours increased the ratio of OR to ORco expression (E). The ratio of the sum of all OR reads in a sample relative to the number of ORco reads in the same sample was significantly lower in the control group compared to con- and allospecific exposure (Wilcoxon tests, * p < 0.05, ** p < 0.01).

To better understand the molecular processes underlying the dynamics of OR gene expression, we analysed the effect of odour exposure on the expression of all genes. Since we were interested in the genes involved in OR profile dynamics independent of colony identity and independent of the specific odour the ants were exposed to, we accounted for colony ID before testing for the overall treatment effect. Out of the 3378 genes expressed in the antennae, 8 were consistently affected by odour exposure across both colonies (LRT padj < 0.05, Tab. S7). Among these genes were orthologues of *histone HSA* variant (Aech_g03759_i1/NP_001262997.1), three genes with products involved in regulating transcription and splicing (*CWC15 homolog, nucleolar protein at 60B, mediator complex subunit 21*), and genes for two proteins involved in neural development (*tubulin-folding cofactor B, tomosyn isoform K*).

Next, we investigated the relative expression of the genes coding for tuned ORx receptor components relative to that of the odorant coreceptor (ORco) to better understand how OR gene expression might relate to the actual OR profiles in the OSN cell membranes. There was a positive correlation between the number of *ORco* reads and the sum of all other *OR* gene reads, as expected when ORx and ORco proteins form functional receptor complexes (p < 0.001, Fig. S1, Tab. S3). Interestingly, the correlation between the number of *ORco* reads and the total *OR* read count differed between treatments (interaction *ORco* reads : treatment p < 0.01, SI Tab. 3). The total number of all *OR* reads combined was higher than the number of *ORco* reads (Fig. 2E). As predicted from the interaction, the experimental odour exposure affected the ratio of *OR* to *ORco* gene expression (Fig. 2E), with higher *OR/ORco* ratios found in the ants exposed to the non-nestmate CHC extracts than in pentane (n = 29, Wilcoxon test control vs. conspecific odour p < 0.01; control vs. allospecific p < 0.05). This was not caused by a change in *ORco* expression (Tab. S6) and might be interpreted as an increase in ORx subunit production after the odour exposure.

## Discussion

The hydrocarbon mixtures covering the cuticles of social insects are colony-specific and form labels that are used for nestmate recognition. We show that the gene expression patterns of the odorant receptors that leaf-cutting ants use for the perception of the labels are also colony-specific, indicating that their profile might be optimised for nestmate recognition. When ants are exposed to non-nestmate labels, they habituate in a way that reduces aggression against the non-nestmate labels (Guerrieri et al., 2009; Stroeymeyt et al., 2010). We show that at the same time, the OR gene expression profiles in the antennae of the treated ants change. This change was odour-specific, as the OR profiles of two colonies exposed to the same non-nestmate colony converged. This indicates that the OR profiles may change dynamically according to the current olfactory environment.

Our findings open up the possibility that the OR profiles are optimised for nestmate recognition. To discriminate nestmates from non-nestmates, certain substances are likely more useful than others: Substances that vary randomly, and those found on nestmates and non-nestmates in similar quantities, are not very informative. Their perception is thus irrelevant for nestmate recognition. In contrast, other substances may differ strongly between friends and enemies, and increasing the sensitivity to those substances would improve nestmate recognition. As has been hypothesized before, changes in OR expression could alter the receptor profiles across the antenna and could optimise recognition (Teşileanu et al., 2019).

One specific case of this optimisation is described as the pre-filter mechanism for nestmate recognition. This hypothesis suggests that neurons that are constantly activated by nestmate-specific substances become desensitized, resulting in the loss of perception of these odorants (Ozaki and Hefetz, 2014). This sensory adaptation should enable individuals to better perceive potential non-nestmate stimuli appearing in their environment. If the production of ORs tuned to nest-specific substances was reduced, this would result in a lowered sensitivity (and lowered reaction to) nest-specific substances. In turn, this downregulation could potentially free resources for increasing the numbers of ORs tuned to substances best suited to perceive novel odours and potentially intruders. Typically, desensitization is expected to occur to receptor complexes (Wicher, 2018) or to neural networks (e.g. in the antennal lobe), but changes in OR numbers across the antenna would have similar effects.

An alternative explanation for our findings would be that the change in OR gene expression is a more or less passive response to OR breakdown caused by OR use. It is plausible that odorant binding to receptors causes wear-and-tear to the receptor complexes. If this causes breakdown of receptor parts, the cell might repair receptors by replacing the entire receptor complex or its parts (Koerte et al., 2018; Von Der Weid et al., 2015). In our experiment, *ORco* expression remained constant, so that it is unlikely that the entire receptor complexes were replaced. While the explanation that ORx subunits are replaced may be parsimonious, some of our observations suggest that there are indeed active changes that would be expected if the OR profiles were constantly optimised for recognition. These clues come from the experimental change of the recognition templates by exposure to non-nestmate labels. Odour exposure did not only trigger changes in OR expression as would be predicted by a passive tracking of damage caused by the most common odorants at any given time. Instead, the exposure also triggered changes in gene expression in the antenna that could indicate that the receptor profiles are actively modified independent of wear and tear of OR complexes. The tubulin-binding cofactor B (orthologue of *Aech_g10112_i1*), for example, is involved in the regulation of neurogenesis in mammals (Lopez‐Fanarraga et al., 2007), and tomosyn (*Aech_g14430_i1*, syntaxin-binding protein 5) is involved in regulating synaptic vesicles so that its knockdown can affect memory formation in *Drosophila* (Chen et al., 2011). We also find genes differentially expressed that are involved in the regulation of transcription or processing of RNA (e.g. splicing) (*Aech_g12595_i1*: mediator complex subunit 21, *Aech_g15049_i1*: nucleolar protein at 60B, and *Aech_g03759_i1*: spliceosome-associated protein CWC15 homolog). A particularly interesting gene is *histone H2A*.*V* (*Aech_g03759_i1*). Its expression is affected by odour exposure as well, hinting at the possibility that histone modification may be a mechanism to regulate colony-specific odorant receptor profiles (Baldi and Becker, 2013).

How exactly the OR profiles are altered is still unclear. Usually, each olfactory sensory neuron (OSN) possesses only one ORx type in the membrane, i.e. one OSN is responsible for the perception of one odorant. OR profile alterations could thus happen through increasing the density of important ORs on the membranes of their respective OSNs, or by replacing the OSNs altogether, which occurs regularly in *Drosophila* and other species (Fernández-Hernández et al., 2020; Lledo and Gheusi, 2005; Yan et al., 2017). In our experiment, the *OR:ORco* gene expression ratio systematically increased after we changed the recognition templates. OR complexes are typically made up by three identical ORx proteins and one odorant coreceptor (Yan, 2024; Zhao et al., 2024). In our control ants, the *OR:ORco* gene expression ratio was around five but the ratio increased in particular in ants exposed to conspecific non-nestmate odour – indicating that more ORx subunits were produced when the label changed. Since the ORx are the receptor parts that are specifically tuned to odorants, one would indeed expect them to be more affected by specific odorant changes than the generic co-receptor ORco in case only the odorant-specific receptor subunits were swapped out. In contrast, the most parsimonious expectation for neuron turnover would be that ORco and ORs would be equally affected. It is thus tempting to speculate that neurons might replace the ORx subunits of the receptor. This would suggest that in case of a systematic change in the olfactory environment, some OR subunits might be swapped out to optimise the sensitivity to important cues, while the co-receptor subunits would be retained.

### Conclusions

No matter the mechanism, we show that OR gene expression in ants is colony-specific and dynamically changing with a changing nestmate template. It is quite possible that the flexibility of OR profiles aids to optimise nestmate recognition, indicating that template formation may start in the antenna, possibly as part of a pre-filter mechanism (Ozaki and Hefetz, 2014). Earlier studies have suggested the involvement of higher brain centres in the formation of the nestmate recognition template (Bos and d’Ettorre, 2012). These processes are by no means mutually exclusive. We suggest that a pre-filter mechanism could act as a secondary mechanism in parallel with learning, or could even be activated by learning. When specific labels are important enough to be learnt, it would be beneficial to be able to accurately identify them. Learning might thus trigger feedback to the antennal lobe and the OSNs to optimise the perception of and discrimination among relevant labels. The nestmate recognition template can be seen not as a single neural representation located in a specific compartment of the neural system, but rather a complex of neural processes from the periphery up to the central nervous system (Sung et al., 2021). The sum of modulations that occur in parallel in central and peripheral olfactory systems could work in synergy and optimise nestmate recognition. This view reconciles different lines of evidence for both peripheral and central nervous system templates (Bey et al., 2025; Stroeymeyt et al., 2010).

## Acknowledgements

We would like to thank the Department of Ecology & Evolution for stimulating discussions and the Global Ant Genomics Alliance for providing us with early access to the revised *Acromyrmex echinatior* genome. We would also like to thank the German Research Foundation (NE 1969/6-1) for funding. Our thanks also go to the Smithsonian Tropical Research Institute in Panama for providing access to their laboratory facilities and to the Autoridad Nacional del Ambiente of Panama for issuing collection and export permits.

## Methods

### Study organism

Twelve queenright *Acromyrmex* colonies collected in Gamboa, Panama between 2017 and 2022 were used for behavioural experiments (9 colonies of *Acromyrmex echinatior*, 3 colonies of *Acromyrmex octospinosus*). Two of the *A. echinatior* colonies were also used for transcriptomic analysis. The ants were maintained at the University of Freiburg at a temperature of 25 – 28°C and 70 - 80% humidity with a 12 h light–dark photoperiod in fluon-coated plastic boxes (from 39 × 28 x14 up to 57 × 39 × 28 cm, depending on the colony size). Each box contained fungus gardens grown in beakers (from 0.5 up to 1L, according to the colony size) covered by upturned plastic flowerpots. The colonies were provided with bramble, rice, and apple slices, and sprayed with water twice a week. The experiments were conducted from December 2021 to February 2023.

### Experimental design

We tested whether an *Acromyrmex echinatior* ant constantly exposed to a new cuticular hydrocarbon profile extract would modulate its behaviour toward that CHC extract, and whether that changed the gene expression in its antennae. We exposed ants in subcolonies to CHC extracts from different colonies belonging to the same or a closely related species. Afterwards, we measured the aggression toward the non-nestmate CHC extract, and the gene expression in the antennae (Fig. 1A). From 7 *Acromyrmex echinatior* original colonies, we set up 42 subcolonies that we exposed to three different exposure treatments. For each subcolony, 8 to 10 medium-sized forager ants were set up in fluon-coated© plastic boxes (10 × 10 cm) and provided with fungus, bramble leaves, and moist cotton (Tab. S4). On the following day, we began to expose each subcolony to either solvent control, conspecific non-nestmate odour, or allospecific non-nestmate odour, for three days. We used glass slides covered with either pentane solvent (control), non-nestmate CHC extract from *A. octospinosus* (allospecific) or from *A. echinatior* (conspecific), and replaced the slides once a day (Fig. 1A). After three days of exposure, we tested whether the ants previously exposed to non-nestmate CHC extract would decrease their aggression towards the CHC extract they had been exposed to. Half of the control ants were tested for their reaction to conspecific CHC extract, and the other half to allospecific CHC extract. For two *A. echinatior* colonies, some of the ants were not used for the behavioural experiment but for the transcriptomic analysis instead (Tab. S4). Their antennae were dissected and the RNA extracted separately for each individual. RNA was sent for sequencing for 28 ants in total.

### CHC extracts

Medium size foraging workers were killed by freezing at −20 °C for 45 minutes, then placed in a glass beaker and covered with n-pentane for three minutes on an orbital shaker (Phero Shaker, Mod.:13A34, Biotec Fischer GmbH) to extract the cuticular hydrocarbons (CHCs) from the ants’ cuticles. The ants were removed using a metal sieve and the solvent was evaporateed. Then, pentane was added to achieve a concentration of 1 ant per 11 μL. The extracts were kept in 5ml glass tubes that were renamed with codes to ensure blind experimentation. We applied 22 μL (2 ant equivalents) of the solution onto glass slides and let them dry before placing them into the subcolonies.

### Behavioural analysis

After 3 days of odour exposure, on day 4, we tested whether the ants would react aggressively to the odour or to solvent control. Behavioural assays were conducted in a circular fluon-coated arena placed on filter paper (Ø 5 cm). A dried slide of 11 μL (1 ant equivalent) of CHC extract was placed inside a smaller plastic cylinder (Ø 2.5 cm) in the middle of the arena and the focal ant was placed outside this cylinder to prevent contact. The ants were left to acclimatize to the arena for five minutes before the small cylinder was removed to allow contact between the individuals and the slide. Tests were live- or video-recorded. The focal ants’ behaviours were quantified using the software Boris (version 7.10.55 (Friard and Gamba, 2016) and the observer was always blind since both the focal ant’s previous exposure and the origin of the CHC extract for the test were unknown. Mandible opening duration near the slide was recorded. Mandible opening is a precursor of biting behaviour and a good estimate of aggression (Guerrieri and d’Ettorre, 2008).

Data points were removed from the dataset for ants that never came into contact with the slide, leaving 232 observations in total (Tab. S4). We had conducted preliminary tests before the experiment to ensure that the focal colonies were aggressive against the colonies used to obtain the CHC extract.

The duration of mandible opening behaviour (continuous variable from 0 to ca. 36 seconds) was set up as the dependent variable in generalized linear models (GLM) with quasi-poisson error family. The different exposure (allospecific or conspecific vs. control), the colony origin of focal ants, and their interaction were used as predictors. The experiments with allo- and conspecific CHC extracts were analysed separately. Significance testing was done using variance analysis tests (anova.glm() function of the stats library). The results were interpreted as significant when p < 0.05 (two-tailed). All statistical analyses were performed using R 4.0.5 software (R Development Core Team, 2016).

### Transcriptome analysis

Antenna tissues were dissected after 3 days of exposure, on day 4. Each sample was composed of both antennas from a single individual. Just after dissection, the antennae were placed in 50 µl peqGOLD Tri Fast™ and crushed with 2 Zirconia beads in a Tissue Lyser II, for 2 minutes at 25 Hz, and where then preserved at −80 °C until extraction. RNA was extracted from the antenna following an optimised RNA isolation protocol using a Qiangen kit, RNeasy©. First, we thawed the tissue for 2 to 3 min and after adding 200 µl peqGOLD Tri Fast™, and homogenized the sample. Then, we added 50 µl of chloroform to separate the aqueous phase, before 100 µl isopropanol was added to precipitate the RNA. After the content was transferred into an RNeasy Mini spin column, we washed the column with the RPE buffer provided by the kit, which we had mixed with ethanol. The sample was centrifuged for 1 min at 4 °C, 11000 xg. The washing step was repeated three times. Finally, the RNA attached to the column membrane was dissolved in 30 µl nuclease-free water.

The RNA samples were kept at −80 °C until they were sent to BGI, Hong Kong. The libraries were prepared by BGI as DNBSEQ Low Input Smart-Seq Eukaryotic mRNA library. Amplified libraries were sequenced on the DNBseq platform (100bp paired-end reads), generating ~8 Gigabases of raw data for each sample.

RNASeq raw reads were trimmed using fastp and filtered using MultiQC (Ewels et al., 2016). The remaining sequences were mapped to the *Acromyrmex echinatior* GAGA genome (Vizueta et al., 2025) using HISAT2. The SAM files thus generated were converted to BAM files and sorted by name using SAMtools. The sorted BAM files were run into HTSeq (Anders et al., 2015) to generate gene count tables using the following settings: -f bam, -i transcript_id, -t CDS, -m union, -r name, -- stranded=no.

We determined the differentially expressed genes (DEGs) between workers exposed to CHCs and pentane (control) using the generalized negative binomial model implemented in DESeq2 (Love et al., 2014). P-values were calculated by Wald tests and corrected for multiple testing using the false discovery rate approach. Genes with adjusted p-values < 0.05 were listed as DEGs. To plot the results, we normalized the gene count data using the rlogTransformation function from DESeq2, which transforms the count data to the log2 scale in a way which minimizes differences between samples for rows with small counts. We performed a principal component analysis (PCA) with the plotPCA function from the DESeq2 package. All statistical analyses were performed using R 4.0.5 software (R Development Core Team, 2016). We used blastn to find *Drosophila* orthologues of non-OR genes of interest.

