## Supplemental Information for "Exposure to non-nestmate odours changes the odorant receptor profile in *Acromyrmex echinatior* ants"

**Supplemental information Bey et al: OR expression and habituation in ants**

**Figures S1–S3, Tables S1-S4**

**Tables S5-S8 in separate files:**

**Table S5.** DESeq2 analysis results for OR comparison between colonies, separately per treatment. Related to Fig. 2A-C.

**Table S6.** Treatment effect on OR expression, separately per colony. Related to Fig. 2D.

**Table S7.** DEGs and significant functional GO terms for the exposure effect, per colony

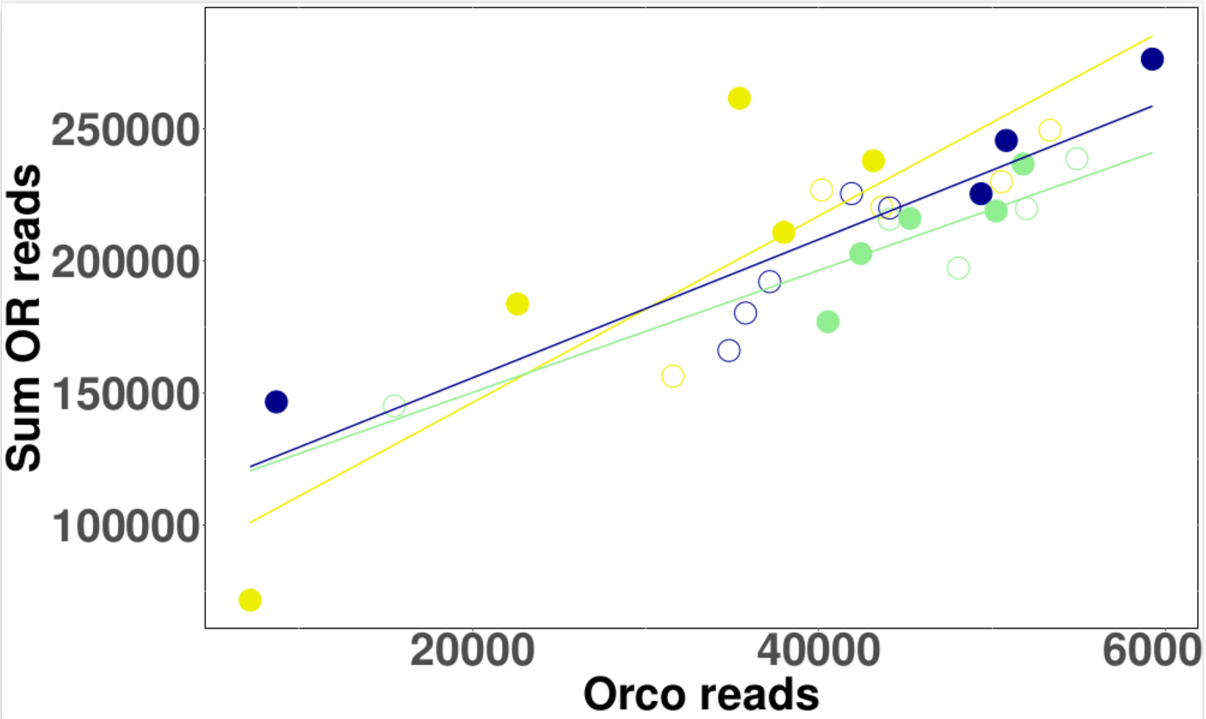


**Figure S1: The total OR gene expression was positively correlated with the expression of the OR coreceptoir orco**. The sum of the OR gene reads depended on the number of ORco reads ($n = 29$ individuals, $p < 0.001$, Tab. S3) and the odour ants have been exposed to ($n = 8$-$10$ per treatment, $p < 0.05$, Tab. S3). Each dot is the measurement of an individual ant's gene expression, the different colours correspond to the different treatments the ants were exposed to. Filled and empty dots indicate individuals from two different colonies.

**Table S1:** Analysis of Variance on a GLM with quasi-poisson error family, describing the duration of mandible opening in reaction to conspecific non-nestmate CHC extract.
Full model: mandible opening ~ Colony origin focal ants x Treatment

Colony origin of focal ants: $F_{6,169}$ = 3.05, p = 0.007
Treatment: $F_{1,168}$ = 0.15, p = 0.697
Colony origin of focal ants x Treatment: $F_{6,162}$ = 2.61, p = 0.016

**Table S2:** Analysis of Variance on a GLM with quasi-poisson error family, describing the duration of mandible opening in reaction to allospecific non-nestmate CHC extract.

Colony origin of focal ants: $F_{3,52}$ = 4.45, p = 0.008
Treatment: $F_{1,51}$ = 11.93, p = 0.001
Colony origin of focal ants x Treatment: $F_{3,48}$ = 0.27, p = 0.845

**Table S3:** Analysis of Variance on a negative binomial glm, n = 29.
Full model: OR read sum ~ Colony ID + Orco reads x Treatment

| | Df | Deviance | Resid. Df | Resid. Dev | p | |
|---|---|---|---|---|---|---|
| NULL | | | 28 | 124.290 | | |
| colony ID | 1 | 0.075 | 27 | 124.215 | 0.784687 | |
| treatment | 2 | 0.128 | 25 | 124.087 | 0.937883 | |
| number of orco reads | 1 | 83.942 | 24 | 40.145 | < 2.2e-16 | *** |
| treatment:orco reads | 2 | 11.081 | 22 | 29.064 | 0.003924 | ** |

**Table S4: List of experimental subcolonies**

| ID | Number of sub-colonies (8-10 ants each) | Number of ants used for the behavioural experiment (ants removed*) | Number of samples for transcriptomic data | Focal colony | Exposure odour | Test odour |
|---|---|---|---|---|---|---|
| 1 | 3 | $n_{Ae}$ = 16 (6) | 5 | Ae21 | Ae34 | Ae34 |
| 2 | 3 | $n_{Ae}$ = 21 (2) | 5 | Ae32 | Ae34 | Ae34 |
| 3 | 1 | $n_{Ae}$ = 5 (4) | | Ae16 | Ae37 | Ae37 |
| 4 | 2 | $n_{Ae}$ = 3 (12) | | Ae7 | Ae40 | Ae40 |
| 5 | 2 | $n_{Ae}$ = 13 (3) | | Ae34 | Ae43 | Ae43 |
| 6 | 2 | $n_{Ae}$ = 15 (1) | | Ae40 | Ae49 | Ae49 |
| 7 | 2 | $n_{Ae}$ = 11 (5) | | Ae43 | Ae49 | Ae49 |
| 8 | 2 | $n_{Ao}$ = 6 | 5 | Ae21 | Ao62 | Ao62 |
| 9 | 1 | $n_{Ao}$ = 4 | 4 | Ae32 | Ao62 | Ao62 |
| 10 | 2 | $n_{Ao}$ = 3 (11) | | Ae21 | Ao2 | Ao2 |
| 11 | 1 | $n_{Ao}$ = 3 (7) | | Ae16 | Ao2 | Ao2 |
| 12 | 2 | $n_{Ao}$ = 8 (7) | | Ae7 | Ao3 | Ao3 |
| 13 | 2 | $n_{Ae}$ = 13 (3) | | Ae34 | Pentan | Ae43 |
| 14 | 2 | $n_{Ae}$ = 14 (2) | | Ae43 | Pentan | Ae49 |
| 15 | 2 | $n_{Ae}$ = 15 (1) | | Ae40 | Pentan | Ae49 |
| 16 | 3 | $n_{Ae}$ = 19 (1)<br>$n_{Ao}$ = 2 (1) | 5 | Ae32 | Pentan | Ao62<br>Ae34 |
| 17 | 5 | $n_{Ae}$ = 17 (5)<br>$n_{Ao}$ = 15 (5) | 4 | Ae21 | Pentan | Ao2<br>Ao62<br>Ae34 |
| 18 | 3 | $n_{Ae}$ = 11 (4)<br>$n_{Ao}$ = 9 (4) | | Ae7 | Pentan | Ao3<br>Ae40 |
| 19 | 2 | $n_{Ae}$ = 3 (4)<br>$n_{Ao}$ = 6(2) | | Ae16 | Pentan | Ao2<br>Ae37 |

* numbers in brackets indicate the number of ants removed from data set because they did not contact the slide with the non-nestmate CHC extract
